# ENRICH: a fast method to improve the quality of flexible macromolecular reconstructions

**DOI:** 10.1101/400317

**Authors:** M. Kazemi, C. O. S. Sorzano, A. Des Georges, J. M. Carazo, J. Vargas

## Abstract

Cryo-electron microscopy using single particle analysis requires the computational averaging of thousands of projection images captured from identical macromolecules. However, macromolecules usually present some degree of flexibility showing different conformations. Computational approaches are then required to classify heterogeneous single particle images into homogeneous sets corresponding to different structural states. Nonetheless, sometimes the attainable resolution of reconstructions obtained from these smaller homogeneous sets is compromised because of reduced number of particles or lack of images at certain macromolecular orientations. In these situations, the current solution to improve map resolution is returning to the electron microscope and collect more data. In this work, we present a fast approach to partially overcome this limitation for heterogeneous data sets. Our method is based on deforming and then moving particles between different conformations using an optical flow approach. Particles are then merged into a unique conformation obtaining reconstructions with improved resolution, contrast and signal-to-noise ratio, then, partially circumventing many issues that impact obtaining high quality reconstructions from small data sets. We present experimental results that show clear improvements in the quality of obtained 3D maps, however, there are also limits to this approach, which we discuss in the manuscript.

## Introduction

Cryogenic electron microscopy (cryo-EM) using single particle analysis is a well-known structural technique to achieve three-dimensional reconstructions of macromolecular complexes in their native state. This imaging modality has recently transitioned between being considered a promising structural technique to become a well-established approach, capable to provide high-resolution structural maps routinely. One of the main advantages of this technique is its ability to visualize macromolecule complexes that would be too large or flexible to be tackled by nuclear magnetic resonance or X-ray crystallography. Biological macromolecular complexes are not rigid entities and usually exist in a range of conformations that play a key role in their function. Macromolecular structures at sufficient resolution and in different physiological conformations provide essential insights into understanding their function and are key to efficiently design new drugs [Boland2017]. However, obtaining high-resolution reconstructions from highly flexible samples remains a challenge. The reconstruction workflow is an inverse ill-posed problem. In cryo-EM of isolated macromolecules, normally referred to as Single Particle Analysis (SPA), macromolecular projection images are obtained without knowledge about the particle’s orientation. This lack of orientation information added to the low signal-to-noise ratios that characterize cryo-EM images imposes computationally demanding reconstruction procedures as marginalized maximum-likelihood based approaches [Scheres2007; Scheres2012a; Lyumkis2013]. High-resolution structure determination requires obtaining homogeneous sets of particles in the same state. Flexible regions of complex will display lower resolutions correlated with the amplitude of movement. In case of macromolecular complexes exhibiting important flexibility or multiple conformations an added level of complication is a drop off in the accuracy of the particle orientation assignment. Thus, particles from a structurally heterogeneous data set must be divided in structurally homogeneous sets or classes. This processing step is called three-dimensional (3D) classification and is usually performed in combination with 3D reconstruction [Scheres2016]. Computational developments in 3D classification methods have made the reconstruction of different conformations at high resolution from heterogeneous cryo-EM data possible [Scheres2007; Scheres2012a; Lyumkis2013; Punjani2017]. The most successful classification method so far is based on an iterative regularized empirical Bayesian strategy [Scheres2012a]. This approach is based on determining the parameters of a model that show the highest probability of being the true one taking into the account the observed data (images) and prior knowledge. Most classification methods divide the data set into discrete groups (classes) with typically different sizes. Homogeneous data sets show a linear relationship between the inverse of the obtained resolution (spatial frequency) and the logarithm of the number of particles employed in the 3D reconstruction [Stagg2014; Stagg2008]. Then, a common problem after 3D classification is that the resolution obtained from the computed homogeneous classes may be limited by the number of particles belonging to these sets. Moreover, there are technical reconstruction issues that become more severe when the number of particles is small, such as unevenness of the angular distribution [Sorzano2001] and even artifacts of the reconstruction method. Indeed, it is common that after 3D classification only one or maybe a few majoritarian classes (corresponding to the most stable states) can be resolved at high-resolution after refinement. In the worst possible scenario, only modest resolution reconstructions (between 9 and 20 Å) will be attainable for all obtained classes. This situation is typically found when the number of particles was low, or the number of selected classes was high. In all these cases, and even assuming that the 3D classification worked correctly, the number of images in these low populated classes represent a technical problem when aiming for high resolution. Modest resolutions between 9 and 20 Å are not sufficient to observe secondary element structures; thus, it is not possible to fully visualize the conformational changes performed by the macromolecule, complicating a molecular understanding of its function. Currently, there is no computational solution to assist users with such issues. The only option users have is returning to the electron microscope again with the hope of collecting enough additional data to resolve the different conformations at sufficient resolution. If resources are available, of course this is the best approach, but in many practical cases it may not be feasible. Recently, it has been proposed solutions to process flexible samples [Nakane2018; Schilbach2017]. In [Nakane2018] is presented an approach that performs independent focused refinements with iteratively signal subtractions of rigid bodies defined by the user. In [Schilbach2017] authors showed the capabilities of a new software called WarpCraft, which uses normal mode analysis to model the motions between different regions of a cryo-EM map.

In this work, we propose a fast and efficient computational approach that can increase the resolution of modestly populated classes without requiring collecting more data, so it is aimed at “making the best” from relatively limited data sets. As an example, our approach required only 2 hours and 50 minutes to process particles from the 80S ribosome presented below using 10 CPUs. The idea has been inspired by classical work in the field of two-dimensional (2D) electron crystallography, where a less-than-perfect 2D crystal is modified (distorted) so that an artificially improved crystal is created [Henderson1986; Gil2006]. The computational method proposed shares some conceptual similarities with the above described 2D crystallography approach. Our method focusses on artificially increasing the number of particles in a class by distorting particles from other similar classes and “moving” them to this class. In this way, as the approach artificially increases the number of images in a small class, then, many technical issues associated with the calculation of a map can be better tolerated, leading to increased resolution reconstructions. However, the big question here is how particles can be “moved” between different conformations, so that, and within certain limits, we can increase the quality of these maps in an objective and quantitative manner.

In this work, we use a regularized optical flow (OF) approach [Bouguet2001] to move (deform) particles between different classes/conformations. We have applied this OF method in the past in the context of frame alignment in cryo-EM [Abrishami2015], where we showed that this algorithm is useful to accurately capture local movements between movie frames. In this case, we use OF with a different goal, which is moving (distorting) particles between two (or more) different conformations (input and reference). In more concrete terms, for a given input particle with orientation Θ and belonging to the input conformation, our approach first computes the 2D OF motion field between the two conformations (input and reference) at orientation Θ. To this end, the map conformations are low-pass filtered and projected at the given orientation Θ, then, the 2D motion field is determined by optical flow approach. Note that map projections are not highly affected by noise, which in principle can introduce difficulties in the robust estimation of the 2D motion field. The obtained motion field is then applied to the input particle which is, therefore, moved from the input conformation to the reference conformation. This process is repeated for all particles belonging to the input conformation. Here, we refer to the reference conformation as the one to be improved and to the input –or inputs–conformations as the ones from which particles are to be moved to the reference one. We envision two clear applications of the proposed approach, both aimed at mitigating map reconstruction problems in classes having a limited number of particles. One is the case when 3D classification results in one majoritarian and one or several additional minoritarian classes. Then, our aim is to push the resolution of the minoritarian classes moving particles from the majoritarian conformation. The other application is based on increasing the number of particles in conformations where all the obtained classes are not highly populated and then the resultant reconstructions are limited by the number of particles. In this case particles from all classes are moved to one reference conformation.

## Methods

### Outline of the method

The goal of this method is to improve the quality of a reference 3D class by moving particles from an input conformation (or conformations) to that reference one; we refer to this approach as enRICH (Resolution Improvement in Conformational Heterogeneity maps). The inputs of the method are therefore the maps representing the different conformations and the particles to move. The particles have to be aligned previously, thus, the method requires the following steps: 1) 3D classification to obtain the different map conformations and the particles associated with them; 2) 3D refinement to obtain accurate particle’s angular assignments; 3) enRICH to move particles from one (or several) input conformations to a reference one; 4) 3D refinement of merged particles belonging to the reference conformation. This pipeline is summarized in Figure 1. In this figure, as an example, our goal is to improve the quality of Class 3 using data from Class 2. The input of enRICH is then the map and particles from Class 2 and the map of Class 3. Therefore, Class 3 is the reference conformation and Class 2 is the input conformation. Steps 1) and 2) from the list above ends with the refinement of particles of each Class. enRICH then starts in Step 3), where we take as input maps from Class 2 and 3 and particles from Class 2. EnRICH produces as output a new set of Class 2 optical flow-deformed particles that we refer to in Figure 1 as “Particles Class2 to Class3”. The workflow continues in Step 4) by merging and refining together corrected particles from Class 2 and particles from Class 3 against Class 3 map, then, obtaining an improved quality map (within certain limits). Observe that enRICH can be used between any classes of Figure 1, but that in this example we have illustrated its use with Class 2 and Class 3 only. Our workflow uses 3D classification and refinement methods from Relion 2.0 electron microscopy package [Kimanius2016], however, any method for 3D classification and refinement can be used as well.

**Figure 1.**
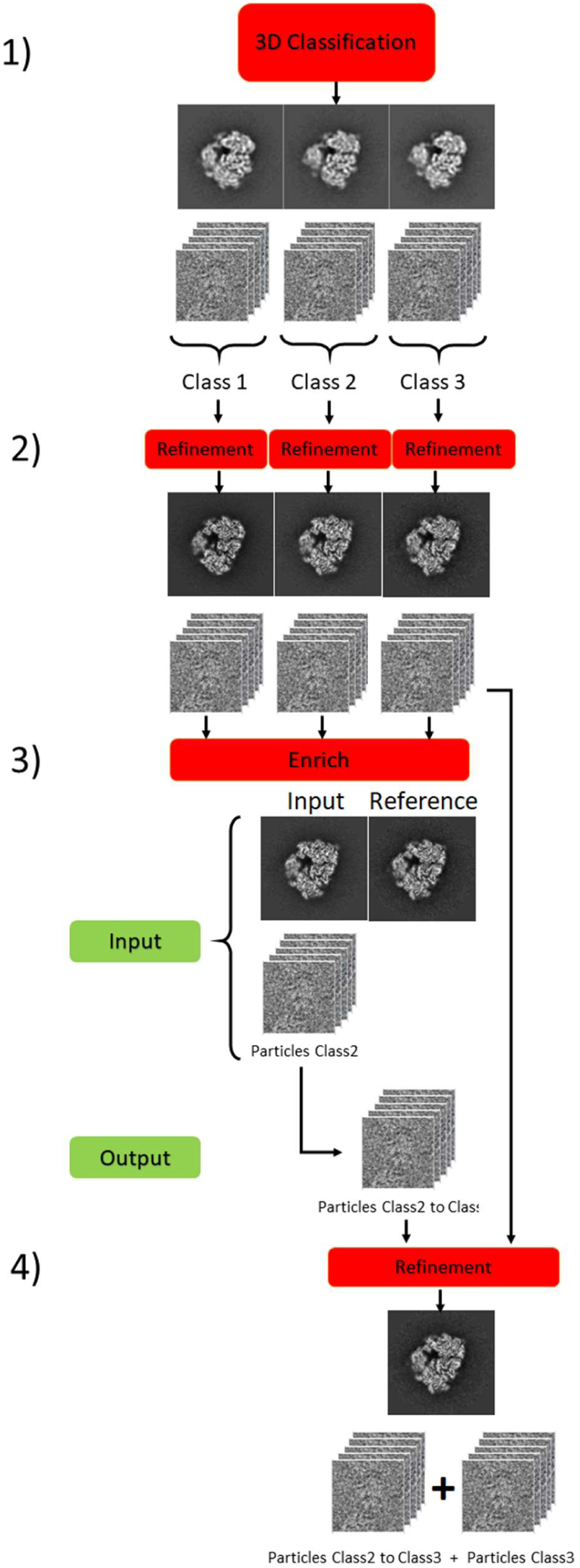
Diagram of the workflow used to improve reconstructions of heterogeneous samples with enRICH.

### Movement of particles between different conformations

In order to move one particle with orientation Θ from conformation A (input) to conformation B (reference), we need to estimate a 2D motion field that maps projected densities from A to B at orientation Θ. The projected densities can be directly obtained projecting aligned maps from conformations A and B at the particle orientation. The 2D motion field can be then determined by optical flow (OF) approach between these two map projections. Here, we use a pyramidal implementation of the Lucas–Kanade OF algorithm with iterative refinement. In the following, we give a summary explanation of this method. For a more detailed description the interested reader is referred to [Abrishami2015; Bouguet2001]. In Figure 2 we show an illustration of the conceptual idea used in Enrich to move particles between conformations.

**Figure 2.**
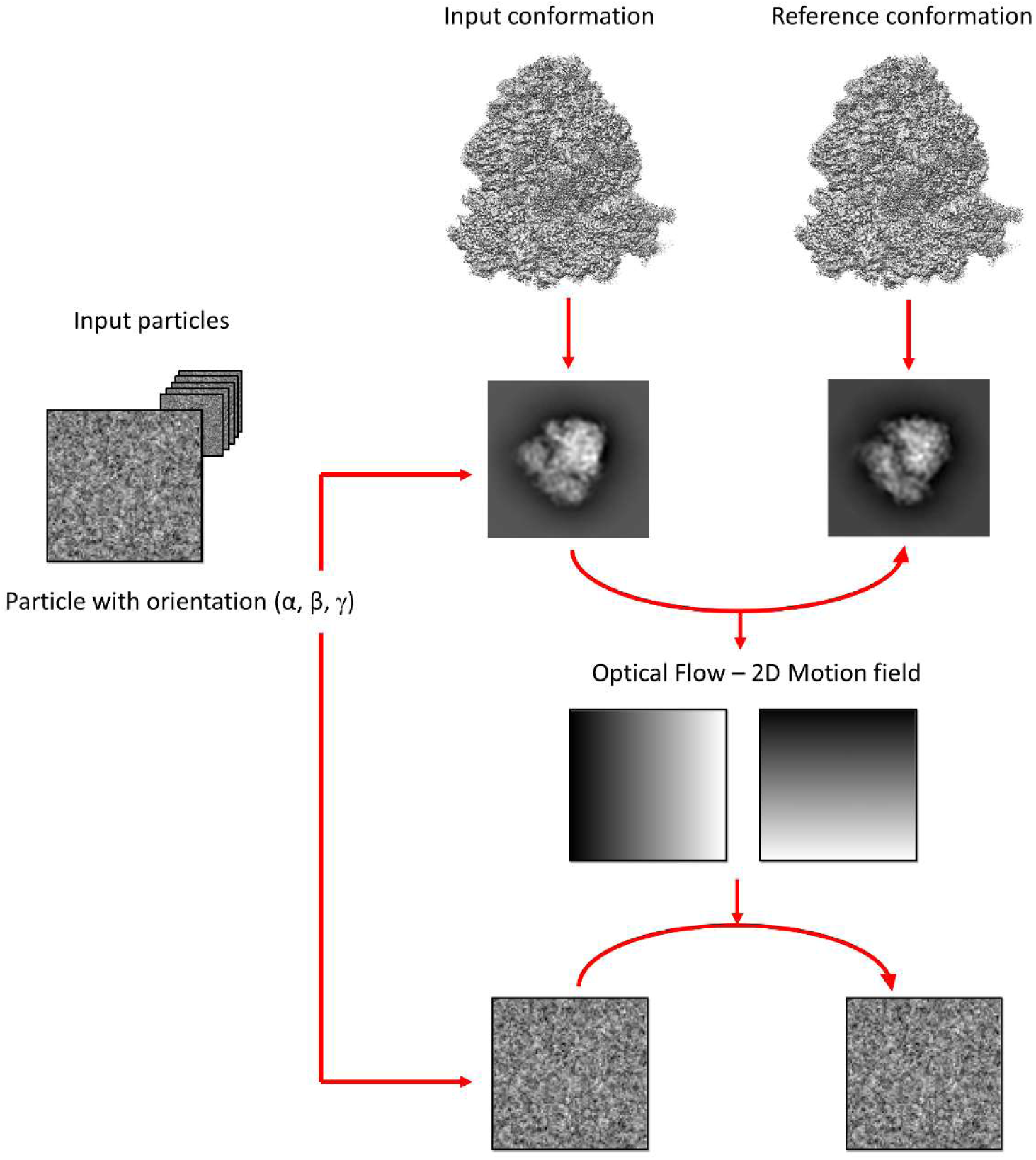
Conceptual idea used in enRICH. The 2D motion (deformation) field between input and reference conformations is obtained by optical flow using map projections at the orientation given by the experimental particle (in this example (α, β, γ)). This particle is then warped from the input conformation to the reference one using this 2D motion field.

Optical flow is a family of algorithms originally introduced in Computer Vision to analyse the movement of objects with respect a common reference frame fixed to the camera. To this end, two consecutive images (*1*) are obtained at times *t* and *t*+ Δ*t* at similar imaging conditions. The intensities of objects within the images at different times do not change and it is their positions that are only modified. Then, the following equation, termed as brightness constancy equation holds

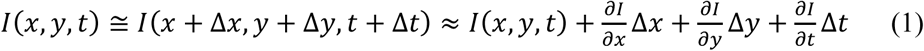

Note that Δ*x* = Δ*x*(*x, y*) and Δ*y* = Δ*y*(*x, y*) refer to the apparent movement of objects between consecutive images. In Eq. (1), we have assumed that the object movements between images are small justifying a first order Taylor expansion. Defining the local shift components *(u, v)* of the 2D motion field as *u* = Δ*x*/Δ*t* and *v* = Δ*y*/Δ*t* we obtain the gradient constrain equation as

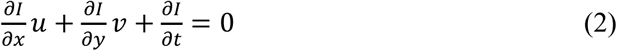

This expression provides one constrain at every pixel, but our goal is to determine two magnitudes *(u, v)*, thus, we have an underdetermined system of equations and is not possible to determine univocally *(u, v)* at every pixel. Lucas-Kanade approach solves this problem imposing the shifts to be similar between close points in the image. Therefore, this approach defines a kernel of size *w × w* around each pixel and imposes the movement to be the same for all pixels inside the kernel, obtaining an overdetermined system of equations. One limitation of the approach outlined above is that it is restricted to cases with small shifts between images that justify the first order Taylor expansion employed in Eq. (1). In cases where this first order approximation is not accurate, it is necessary to iterate multiple times on this scheme. Therefore, after the k^th^ iteration, the brightness constancy equation corresponds to

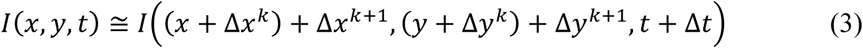

The final displacement vectors obtained after K iterations are as

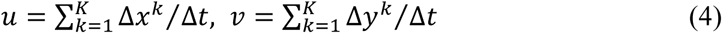

A key parameter of the LK algorithm is the kernel size. This kernel introduces a trade-off between accuracy and robustness of the approach. Large kernels give more robustness to noise and outliers, but less accuracy. Additionally, large kernels are required to handle large motions between images [Bouguet2001]. To solve these limitations, in [Bouguet2001] it was introduced a pyramidal implementation of the iterative LK algorithm presented above. This method is based on performing the recursive LK optical flow approach outlined above over different resolution representations of the input images, called pyramids representations, obtained by down sampling of the input images. The approach performs then by applying optical flow starting with the images with lowest resolution and proceeding through the ones with highest resolution. This process provides coarse estimation of the motion vectors from low resolution pyramid representations that are refined in the next optical flow estimations using higher resolution pyramidal representation images.

### Avoiding model bias in Enrich

The main risk of using OF to move particles between different conformations is the possibility to produce model bias. To avoid this undesired situation, we perform this task in a controlled way. First, the 2D motion fields are calculated by OF without “seeing” the experimental particles to modify. These fields are obtained from filtered map projections at the orientations given by the single particles. The maps are filtered at 15-20 Å resolutions before projection so only large conformational changes will be detected by OF. Additionally, we also use large integrating kernels in OF calculations, of typically between 20×20 or 50×50 Å, that give strong robustness against noise and to prevent overfitting in the determination of the 2D motion fields.

In addition to these two strategies to evade model bias, we have also implemented enRICH following a Gold-standard approach [Scheres2012b]. In this case, the approach uses as inputs half maps at each conformation (input and reference), so that we have a “Half 1” and “Half 2” data sets (maps and particles) in each conformation (input and reference) and the two halves are treated independently.

### Limitations of the method

The working hypothesis of the approach is that large regions of the macromolecule (approximately of ∼15-20Å) move collectively, imposing a strong spatial coherence in the motion field between conformations. Therefore, the present method can be used to capture movements of large regions as subunits, but not to determine local patterns of flexibility of smaller elements, such as secondary structures; additionally, it may introduce geometrical distortions not compatible with good stereochemistry parameters. Consequently, this approach is meant for those cases of collective movements where the 2D motion field between conformations is smooth, continuous and obtainable from map projection images. In other words, when conformational changes are not very large and when there is an element of continuous flexibility underlying structural variability. These assumptions require that 3D movements between conformations be small enough and typically limit the attainable resolution to 4-5 Å. However, in the results section, we also show an example of a structure that is improved beyond 4 Å, corresponding to a case in which the hypothesis of smooth, continuous structural variations holds particularly well. However, in general, it is very difficult to give a clear answer about the range of movements that are within reach of this approach, but we have always found that incorrect application of this approach to structures with large conformational changes can be easily detected *a posteriori*. Indeed, these cases show a decrease in the obtained resolution after particle merging when this value is compared to the resolution obtained before particle merging.

## Results

We have used the proposed method with data coming from the *Plasmodium falciparum* 80S ribosome, EMPIAR ID 10028 [Wong2014], and from the type-1 ryanodine receptor (RyR1) [DesGeorges2016].

### Case with moderate flexibility: 80S Ribosome data set

Ribosomal particles were obtained with a FEI Polara 300 microscope equipped with a Falcon II camera. The number of projection images deposited is 105,145. We first classified the data set in three classes using Relion through Scipion framework [DelaRosa2016]. Class 1 (25,323 particles) was composed by particles without 40S subunit, so these particles were not taken into consideration in subsequent analyses. The rest of particles (79,822 particles), distributed in Class 2 (45,059 particles) and Class 3 (34,763 particles), were further refined independently obtaining reconstructions of 4.43 Å and 4.68 Å, which were improved to 3.45 Å and 3.63 Å respectively after postprocessing (masking and B-factor correction). We then used enRICH to improve first Class 2 and then Class 3 reconstructions. The processing times were 2 hours and 50 minutes and 6 hours to deform the sets of 34,763 and 45,059 particles using 10 and 5 CPUs respectively. After refinement of merged particles, we obtained resolutions of 4.05 Å and 4.05 Å (without postprocessing), and of 3.24 Å and 3.26 Å (with postprocessing) for Class 2 and 3 respectively. The corresponding FSCs of these results are shown in Figure 3(a) and (b). As can be seen from these figures there is a substantial improvement in the resolution for both Class 2 and Class 3 after running enRICH. In all cases, the obtained FSCs using enRICH surpass the ones attained without using enRICH. In Figure 3(c-f) we show map slides of reconstructions obtained for Class 2 with (c) and without using enRICH (d) and for Class 3 with (e) and without using enRICH (f). Figures 3(c-f) are shown without any postprocessing. Figure 3(g) shows 3D maps of the “head” of the smaller ribosomal subunit obtained with enRICH (left) and with the consensus reconstruction using the complete set of 79,822 particles but without any correction (right). This region corresponds to the more flexible part of the structure and is marked with a red rectangle in Figure 3(c). These maps were masked by the same soft-mask and are shown at the same density threshold value. As can be seen from Figure 3(g) the map obtained with enRICH shows better structural details and the densities are not broken as happen in the other case. Notably, the B-factor applied to the map corrected by enRICH was higher (−131.445) that in the other case (−129.146).

**Figure 3.**
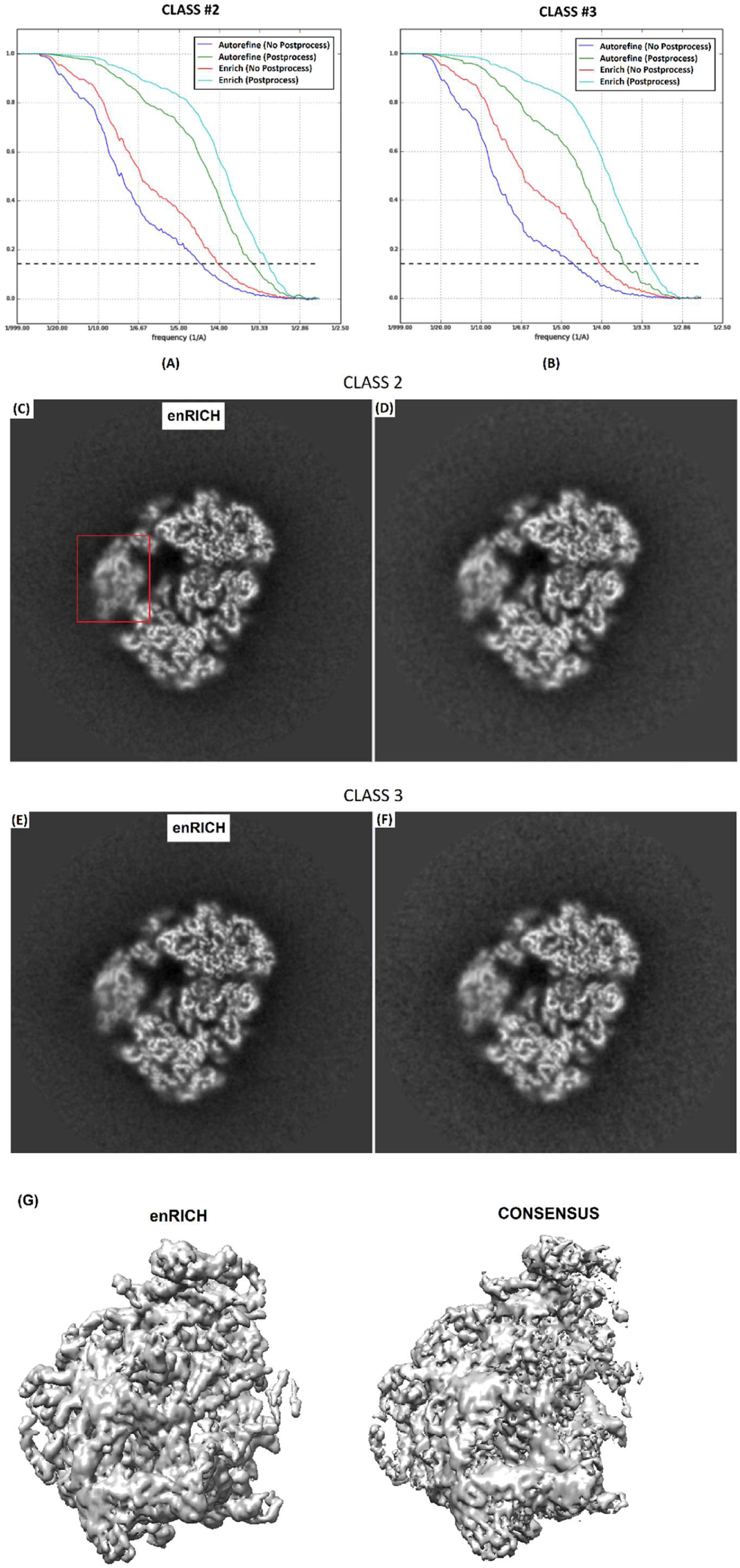
(a-b) obtained FSCs for Class 2 (a) and Class 3 (b) after refinement for particles belonging to these classes (blue and green lines) and for the complete data set after correction by enRICH (red and cyan). Blue and green curves show results without applying any postprocessing, while red and cyan curves present results after masking and performing B-factor correction to reconstructions. (c-f) respective map slides of reconstructions obtained for Class 2 with (c) and without using enRICH (d) and for Class 3 with (e) and without using enRICH (f). The red mark corresponds to the “head” region of the smaller subunit. In (g) 3D reconstructions of the “head” region are shown after correction with enRICH (left) and without any correction by enRICH and using the complete data set (consensus reconstruction). In both cases the number of particles is 79,822.

We could not find differences between the maps obtained with enRICH using and without using the gold-standard approach. In this context, enRICH gold-standard means that every particle is deformed using the 2D motion field calculated from input and reference maps having the same half_id value than this particle, while non using gold-standard in enRICH just means that all particles were processed together using only one map as reference and another one as input. Therefore, particles with half_id=1 (or 2) are deformed by the OF 2D motion field using map projections from maps with half_id=1 (or 2) exclusively. This result shows that enRICH is not affected by overfitting. In Figure 4 we compare FSCs of reconstructions obtained with enRICH using and not using a gold-standard approach for Classes 2 and 3. Figure 4 shows no essential differences between reconstructions in both cases. As can be seen from Figure 3(a-b), the FSCs obtained after Relion auto-refine without postprocessing (blue lines) drops off to ∼0.9-0.8 in the rage between 10-20 Å. These small differences between maps can be propagated to the slightly differences in the FSCs shown in Figure 4. The different 0.143-FSC resolution results are summarized in Table 1.

**Table 1.** Summary of 0.143-FSC resolutions obtained for Class 2 and Class 3 reconstructions and shown in Figures 4 and 5.

|  | **Class #2** | **Class #3** |
| --- | --- | --- |
| **No enRICH + No Postprocessing** | 4.43 | 4.68 |
| **No enRICH + Postprocessing** | 3.45 | 3.63 |
| **enRICH + No Postprocessing** | 4.05 | 4.05 |
| **enRICH + Postprocessing** | 3.24 | 3.26 |
| **enRICH (Gold Standard) + No Postprocessing** | 4.06 | 4.07 |
| **enRICH (Gold Standard) + Postprocessing** | 3.26 | 3.26 |

**Figure 4.**
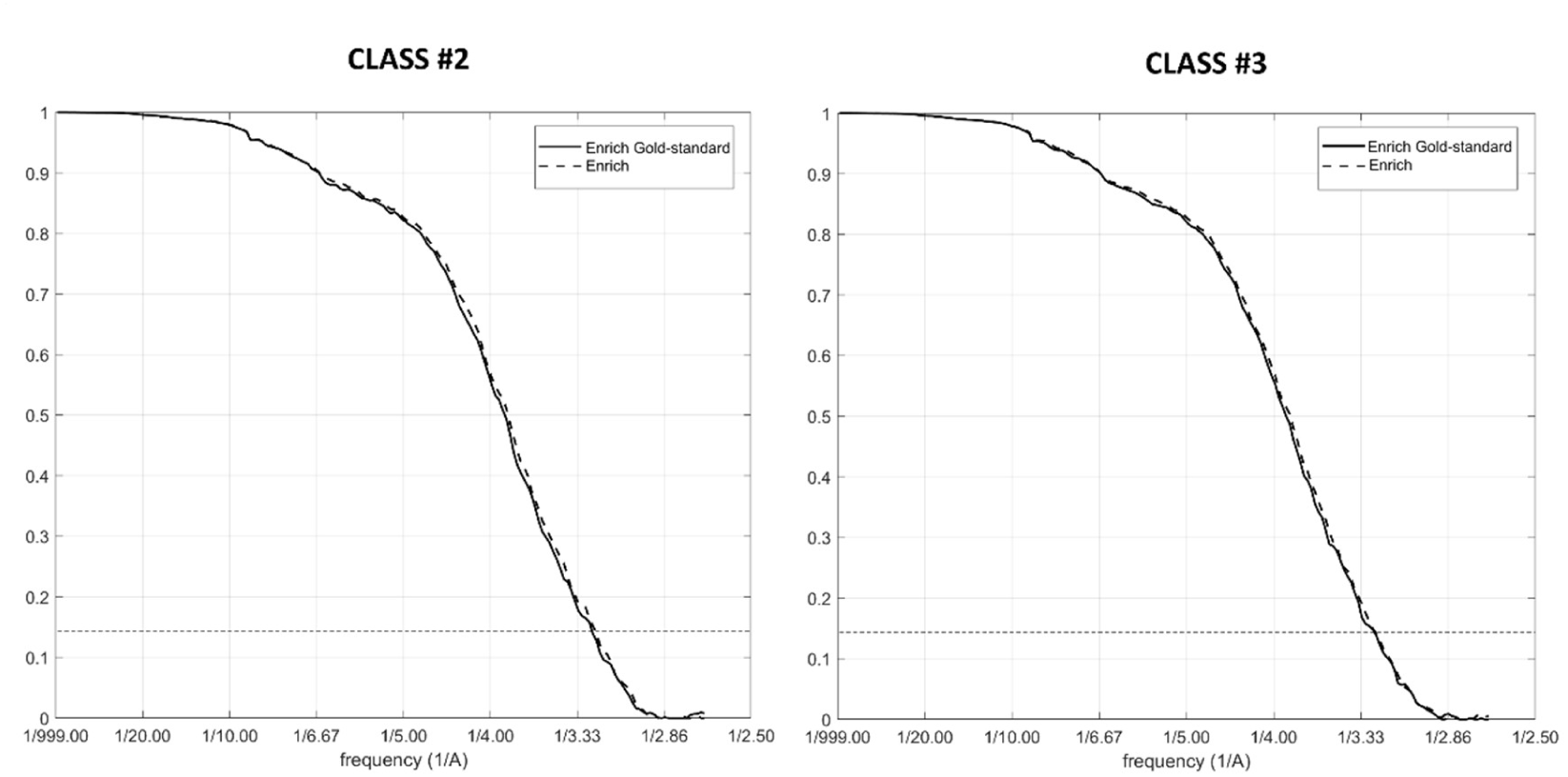
FSCs of reconstructions obtained after refinement and particle correction with enRICH following and not following a gold-standard approach for Class 2 and Class 3.

We compared also the obtained maps with the corresponding PDBs (PDB codes: 3j79/3j7a), since it was critical to show that at this relatively high-resolution reconstruction, the improved maps still conserved coherent structural information. PDB 3j79 corresponds to the bigger 60S subunit, while PDB 3j7a refers to the smaller 40S one. Figure 5 shows the obtained results for Class 2 and 3 using (solid line) and not using (dash line) enRICH approach. The obtained FSC resolutions are presented in Table 2. As can be seen from Figure 5, enRICH improves the different FSC curves for both classes at almost all frequencies.

**Table 2.** 0.143-FSC resolution values obtained for Class 2 and 3 using and not using enRICH approach when reconstructed maps were compared with corresponding PDBs. In all cases, the maps were masked by the same mask and b-factor corrected using similar parameters.

|  | **Class #2** | **Class #3** |
| --- | --- | --- |
| **No enRICH – 3j7a** | 3.40 | 3.56 |
| **No enRICH – 3j79** | 3.17 | 3.34 |
| **enRICH – 3j7a** | 3.22 | 3.23 |
| **enRICH – 3j79** | 3.05 | 3.06 |

**Figure 5.**
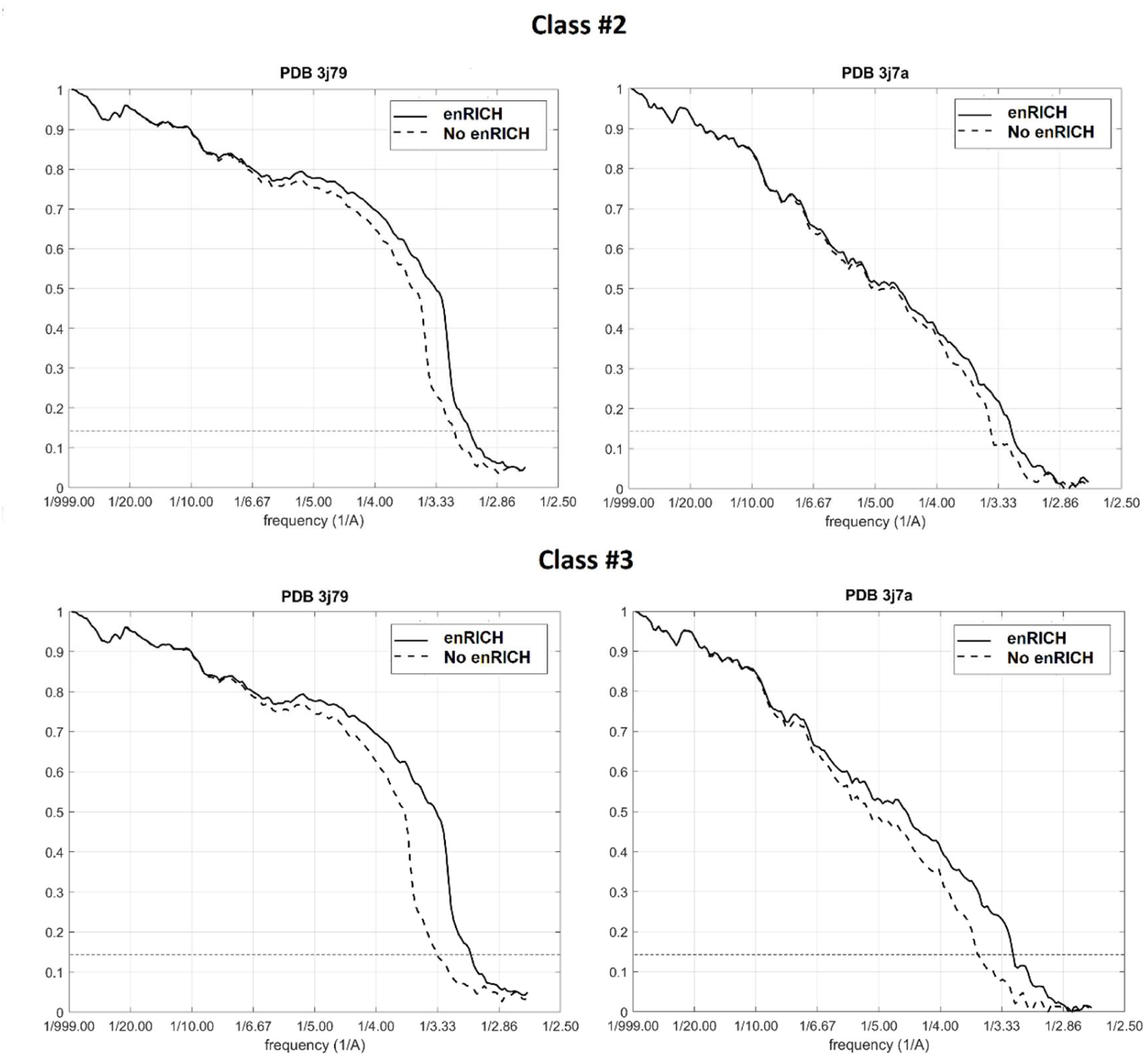
FSC curves obtained confronting corresponding PDBs with maps obtained when only particles belonging to Class 2 and 3 (dashed curve) were used, and when we employed all the particles after correction by enRICH (solid line).

Finally, to show the capacity of the approach for classes with very low number of particles, we further classified the data set composed by merged particles (without any correction) from Classes 2 and 3, into 4 classes. The population of each class was 25,087, 23,702, 5,744 and 24,289 respectively. These classes refined to resolutions of 5.30, 5.74, 15.07 and 5.81 Å without any applied mask or B-factor correction. We then applied enRICH to move particles from Class1, 2 and 4 to Class 3, which is the least populated class. Figure 6 shows the gold-standard FSCs obtained after 3D refinement by Relion auto-refine, without masking or performing B-factor correction for Classes 1, 2, 3 and 4 and after merging all particles to Class 3 using enRICH (blue curve). As can be seen from Figure 6, the resolution is substantially improved after using enRICH.

**Figure 6.**
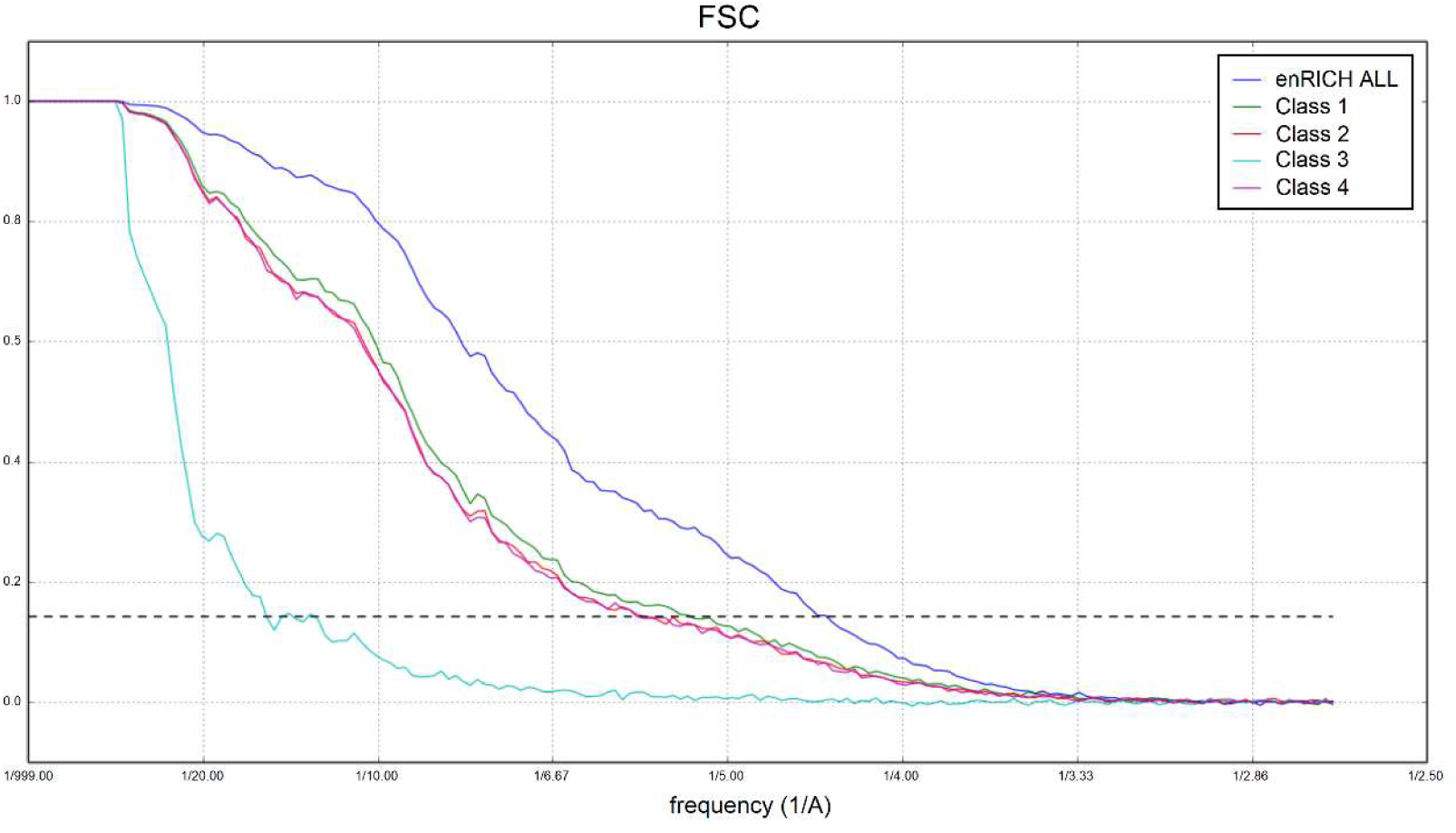
FSCs obtained for Class 1, 2, 3 and 4 reconstructions (green, red, cyan and pink) and after merging all particles to Class 3 using enRICH (blue line).

### Case with large flexibility: RyR1 data set

This data set was composed by 277,044 single particles of RyR1 bound to Ca2+, ATP and caffeine. We first classified this data set into two sets composed by 129,873 (Class 1) and 147,171 (Class 2) particles respectively. Class 1 and Class 2 refined until 4.85 Å and 4.90 Å (0.143-FSC gold standard) using Relion. In this case, we found that the large conformational changes shown by the RyR1 prevented our method to improve these results. Indeed, after moving particles with enRICH from Class 2 to Class 1 and refine the merged set, the resolution achieved was 4.95 Å (0.143-FSC gold standard). As mentioned earlier, complexes with large conformation movements as the RyR1 specimen, prevent our method to obtain high-resolution reconstructions. In these situations, our approach should be used to improve maps at intermediate resolutions only. Therefore, in this case it is not realistic the use of enRICH for improving map resolutions better than 5 Å. However, although we could not improve these results, we employed this data set to show a more appropriate use of our approach, where enRICH was able to improve intermediate resolution reconstructions coming from low populated classes.

In the second experiment, we decided to further classify Class 2 (147,171 particles) into four classes. The particle distribution between classes and the obtained resolution after refinement were Class 2.1: 38,543 particles, resolution 7.17 Å; Class 2.2: 39,239 particles, resolution 7.17 Å; Class 2.3: 35,174 particles, resolution 7.28 Å; Class 2.4: 34,215 particles, resolution 7.28 Å. We then moved particles from Class 2.2 to Class 2.1, particles from Class 2.2 and from Class 2.3 to Class 2.1, and particles from Class 2.2, Class 2.3 and Class 2.4 to Class 2.1. The processing time to deform ∼40,000 particles of 400×400 pixels using 44 CPUs was 1 hour and 35 minutes. In each case, we refined the merged sets obtaining respective FSCs from Relion, that are shown in Figure 7(a). As can be seen from Figure 7(a), the resolution of Class 2.1 was improved from 7.17 (38,543 particles) to 5.35 Å when the dataset was composed by 147,171 particles. Additionally, in Figure 7(b-c) we show respective map slides (without any postprocessing) obtained after merging all particles by enRICH and refining with Relion (Figure 7(b)), and after refining the data set of 38,543 particles alone (Figure 7(c)). As can be seen from these figures, the map slide obtained after merging all particles with enRICH shows better resolution.

**Figure 7.**
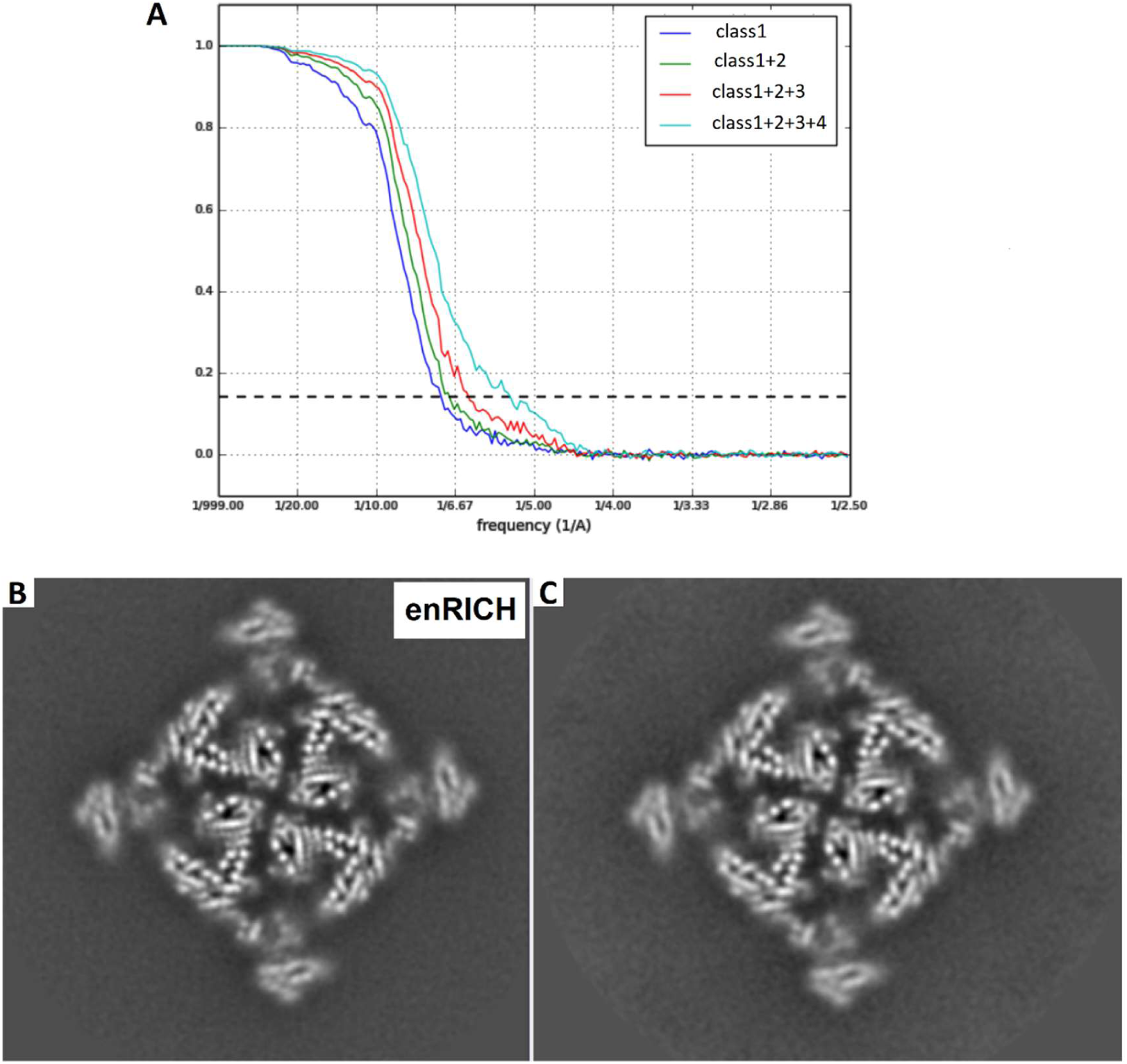
(a) Gold-standard FSCs obtained from Class 2.1 particles (blue curve), Class 2.1 + Class 2.2 (corrected) (green curve), Class 2.1 + Class 2.2 (corrected) + Class 2.3 (corrected) (red curve), Class 2.1 + Class 2.2 (corrected) + Class 2.3 (corrected) + Class 2.4 (corrected) (cyan curve). (b-c) respective slides of maps obtained after merging all particles by enRICH and then refining with Relion (b), and after refining Class 2.1 data set (38,543 particles) alone (c).

The third case we show in this work is aimed at clearly making the point of the trade-off presented by our method between extend of structural variations it can accommodate and the achievable improvement in resolution. In this way, in the third experiment we decided to show that our method can improve maps at moderate resolutions in the presence of large conformational heterogeneity. To this end, we obtained two subsets from Class 1 and two more from Class 2 composed by 10,000 and 5,000 particles randomly selected. Then, we refined the 10,000 particle sets with Relion obtaining resolutions of 8.10 Å for both Class 1 and Class 2. After that, we refined the set of 5,000 particles from Class 1 providing a reconstruction of 12.97 Å. We then used enRICH to move the 5,000 particles set from Class 2 to Class 1, and we merged this corrected (deformed) set with the one composed by 5,000 particles originally coming from Class 1 (and, therefore, without any correction). This set of 10,000 particles (after enRICH correction) reached 8.10 Å (0.143-FSC gold standard) after refinement with Relion. The different obtained FSCs are shown in Figure 8(a). These results show that a reconstruction with 5,000 particles from Class 1 could be improved with enRICH moving 5,000 particles from Class 2. The improvement in map resolution was from 12.97 Å to 8.10 Å.

**Figure 8.**
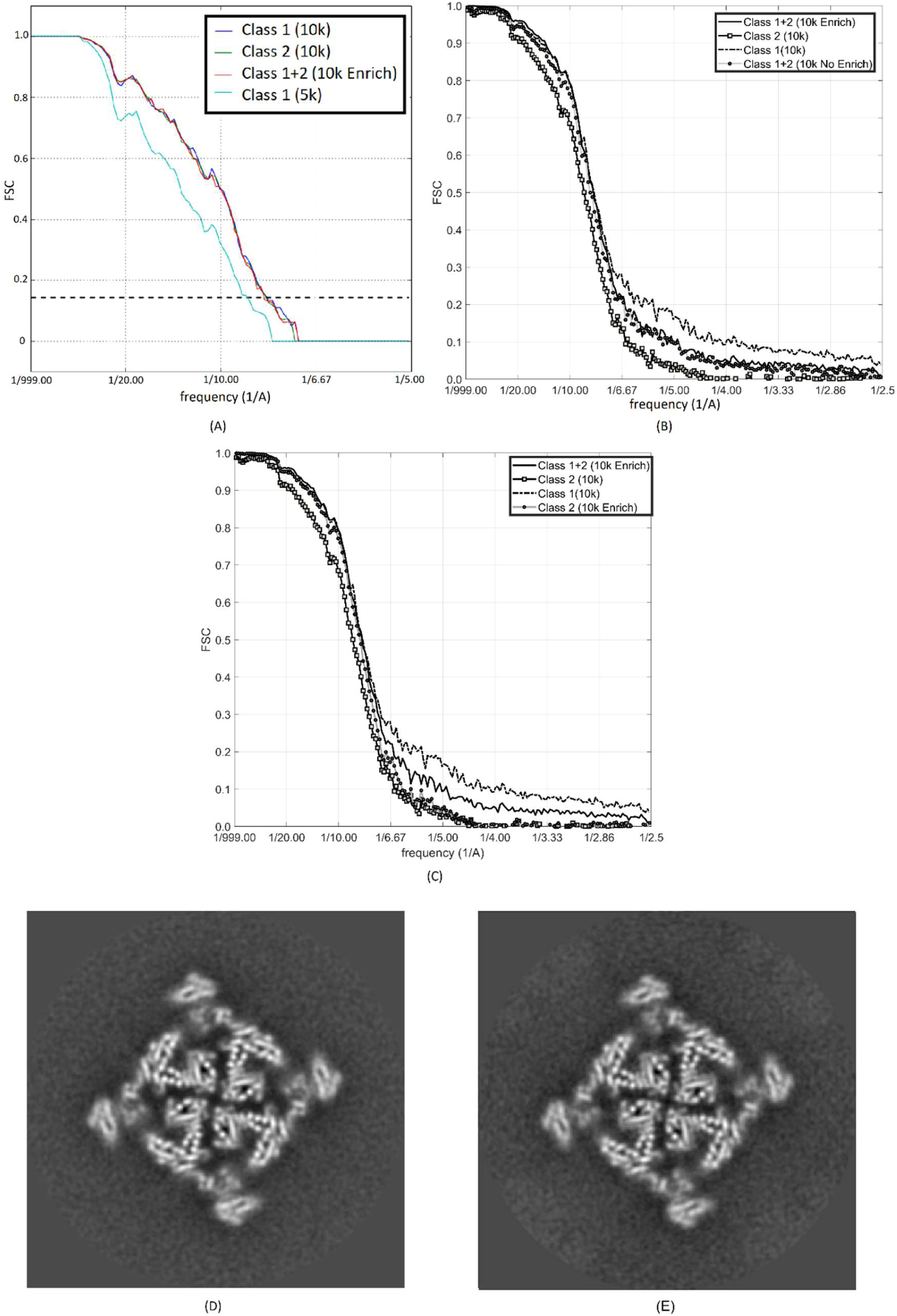
(a) Gold-standard FSCs obtained from 10k extracted randomly from Class 1 (blue line), 10k particles extracted randomly from Class 2 (green line), 5k particles extracted randomly from Class 1 (cyan line) and 10k particles composed by 5k particles from Class 1 and 5k particles from Class 2 moved to Class 1 by enRICH. (b) FSCs obtained comparing the map reconstructed using the complete set of 129,873 particles of Class 1 (ground truth) with maps reconstructed from 10k particles extracted randomly from Class1 (dashed black curve) and Class 2 (solid line with squares) and from merged particles using 5k particles from Class 1 + 5k particles from Class 2 without any correction (grey curve with circles) and after correction using enRICH (solid black line). In (c) are shown similar FSC curves that in (b) but in this case the grey curve with circles corresponds to the FSC obtained from 10k particles from Class 2 corrected to Class 1 by enRICH. (c-d) show respective slides of reconstructed maps from 10k particles of Class 1 and 10k particles composed by 5k of Class 1 and 5k of Class 2 moved to Class 1 after correction by enRICH.

We then compared through FSC analysis these different reconstructions with the map obtained from the complete image set of Class 1 (129,873 particles), which refined until 4.85 Å, and was used as our “ground truth” for structural comparisons. In Figure 8(b), we show the different obtained FSC curves. For the sake of clarity, we also include the comparison between the “ground-truth” and the map obtained after merging the 5k Class 1 particles with the 5k Class 2 particles but without any correction. Figure 8(b) shows the similarity of our “ground truth” map with reconstructions obtained from: (i) 10k particles of Class1 (dashed curve) and (ii) 5k particles from Class 1 + 5k particles from Class 2 corrected with enRICH (solid curve). Curves from enRICH and Class 1 particles are virtually the same until a resolution of ∼6 Å, while FSC curves from Class 2 and merged particles without correction by enRICH present lower FSC values at all frequencies. Beyond this resolution (∼6 Å), FSC curves from merged (with and without correction) and Class 1 particles come near zero, but they do not reach this value. At these resolutions, the FSC values for Class 1 particles are approximately two times the respective ones for the merged particles curves. This behaviour shows that there are correlations between our “ground truth” map and these maps at these resolutions. These correlations are caused because the 10k particles of Class 1 (dashed line Figure 8(b)) and the 5k particles of Class 1 used in the merged reconstructions are contained in the set of 129,873 particles used to reconstruct our Gold-standard. Figure 8(b) shows that enRICH corrects structural differences between different conformations (Class 1 and Class 2 in this example). Figure 8(c) shows similar FSC curves that Figure 8(b), however, in this case, we compare maps reconstructed from 10k particles of Class1 (dashed curve), 10k particles of Class2 (solid curve with squares) and the same 10k particles of Class2 but corrected by enRICH to Class 1 (grey curve with circles) with the “ground truth”. The set of 10k particles of Class 2 corrected by enRICH refined to 8.10 Å using Relion. Again, Figure 8(c) evidences the structural corrections induced by enRICH. Finally, Figure 8(d) and (e) shows similar map slides of reconstructed maps from 10,000 particles of Class 1 and 10,000 composed by 5,000 Class 1 and 5,000 Class 2 after correction by enRICH. These figures show that these sections are very similar.

## Discussion

Macromolecules are not static entities and typically exhibit different conformations when performing their function. Cryo-EM is a high-resolution structure determination technique that can capture these different states providing essential information about the macromolecule’s function. Hence, 3D classification is a routine and crucial task when processing cryo-EM data. However, classifying data requires dividing the collected data into sets of homogeneous particles, a process that usually limits the attainable resolution for the different conformations. These situation is more dramatic in complexes showing a large (or infinite) number of different possible conformations [Sorzano2016]. In this work, we propose a method to merge particles from different conformations. This approach can be used in cases where the resultant number of particles after 3D classification limits the attainable resolution. Then, this method can be used to improve (within limits) the quality of reconstructions coming from minoritarian classes, for example. The approach is based on obtaining the 2D motion or deformation field by a regularized optical flow approach. As shown from our results the approach is fast and computationally efficient. The OF is computed between projections of different conformations at certain orientations given by the experimental particles to process. These motion fields are then applied to corresponding particle images to move them between conformations. The conceptual idea of the proposed method is close to crystal unbending implemented in 2D electron crystallography [Henderson 1985] that enabled a dramatic improvement in resolution. The main goal of crystal unbending is to correct the crystal distortions, that is, its deviation from regular lattice repetition. This technique first obtains the 2D motion field of unit cells with respect to their perfect regular positions and then correct these shifts by interpolation obtaining a corrected crystal image. In both 2D crystallography and single particle analysis underlying physical limitations are crystal local deformations in the first case and large-scale low frequencies motions in the second.

We have taken special attention in avoiding problems related with model-bias when using our approach. First, we use a robust optical flow approach which imposes strong regularization in the obtained 2D motion fields. Typical integration kernels and imposed resolutions to map projections used in the estimation of the motion fields by OF are of 25-50 Å and 15-20 Å, respectively. Additionally, enRICH can be used in a gold-standard fashion. In this case the method uses half maps of each conformation (input and reference) to determine motion fields. Particles are corrected according to their corresponding halves.

It is important to mention that our method imposes a deformation model. This model assumes that the 2D motion field between conformations is smooth, continuous and obtainable from map projection images. These assumptions are only fulfilled when the 3D movements between conformations are not very large and, better still, when there is an intrinsic element of continuous flexibility in the macromolecule under study. Our approach is addressed to capture and correct movements of large regions as subunits, but not to determine small local changes of smaller elements such as secondary structures.

We have applied our approach to two different heterogeneous samples: 80S ribosome and RyR1. In the case of the 80S ribosome, we clearly improved resolution results obtained using pure classes after 3D classification. This result shows that the deformation pattern followed by these ribosomal particles aligns well with our model. On the other hand, we also showed a case where enRICH could not improve resolutions of homogeneous sets after classification. In this situation our deformation model may not describe well the large movements exhibited by the RyR1 macromolecules at high-resolution, limiting the attainable resolution to ∼5 Å. However, we employed this data set to show other practical uses of our approach. In these cases, we clearly showed an improvement in the map resolution after using enRICH. Furthermore, we showed that the approach had not introduced any unwanted bias or artefact and that the resulting (merged) reconstruction was more similar to the original class when reconstructed with a very large number of images that before the merging.

## Acknowledgments

This work was supported McGill start-up funds and by grants from the Natural Sciences and Engineering Research Council of Canada – NSERC (RGPIN-2018-04813), Comunidad de Madrid (S2017/BMD-3817) and the MINECO – Spain (BIO2016-76400-R(AEI/FEDER, UE)).

